## Supplementary figures and tables for "How long does it take to cue a *Pseudomonas aeruginosa* chronic infection phenotype in the lab? Insights from transcriptome analysis in a cystic fibrosis lung model"

***Pseudomonas aeruginosa* transcriptome analysis in a cystic fibrosis lung model reveals metabolic changes accompanying biofilm maturation and increased antibiotic tolerance over time.**

**Niamh E. Harrington^1,2*^, Freya Allen^1,3^, Ramón Garcia-Maset^4^, Freya Harrison^1*^**

1: School of Life Sciences, Gibbet Hill Campus, The University of Warwick, Coventry, CV4 7AL, UK.

2: Department of Evolution, Ecology and Behaviour, Institute of Infection, Veterinary and Ecological Sciences, University of Liverpool, Liverpool, L69 7ZB, UK.

3: (Current address) Institute of Microbiology and Infection and Department of Microbes, Infection and Microbiomes, School of Infection, Inflammation and Immunology, College of Medicine and Health, University of Birmingham, Birmingham, B15 2TT, UK.

4: Warwick Medical School, Gibbet Hill Campus, The University of Warwick, Coventry, CV4 7AL, UK.

*SUPPLEMENTARY FIGURE AND TABLES*

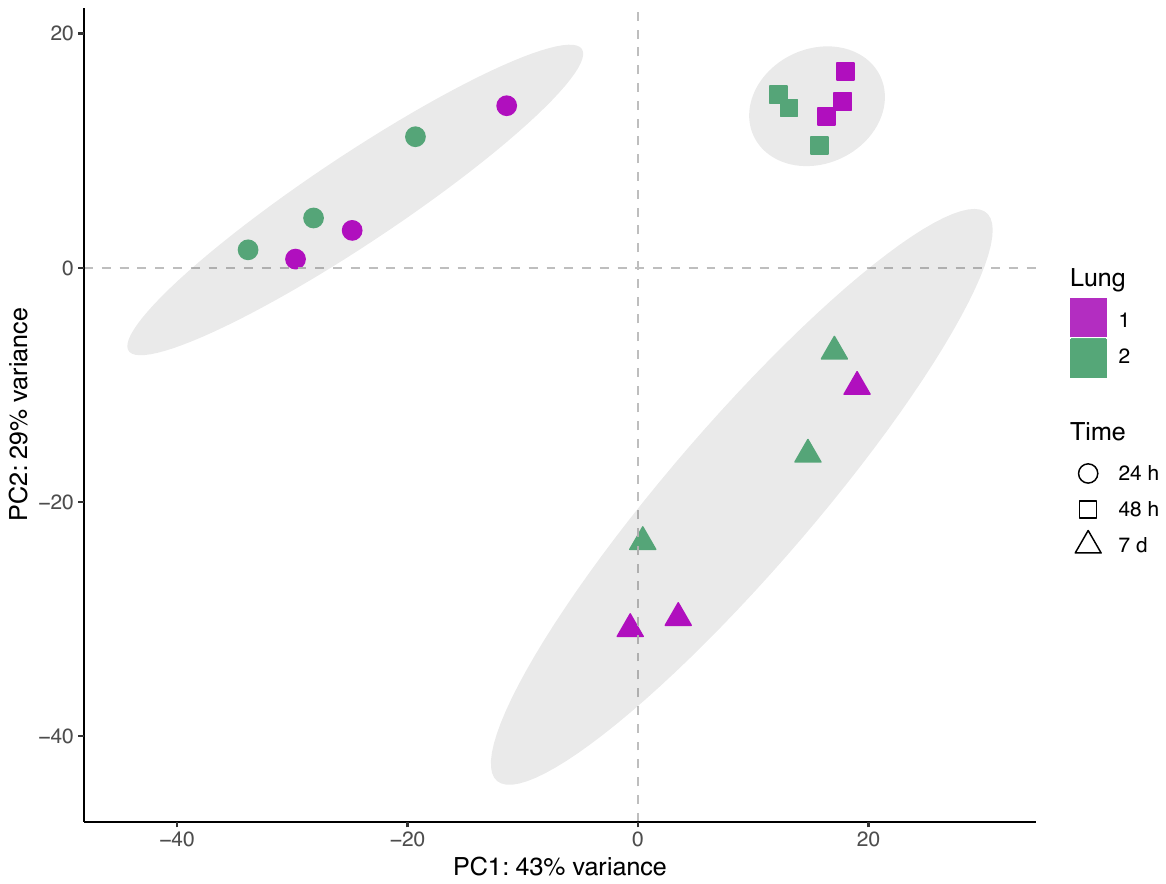
**Figure S1. Principal component analysis (PCA) plot of the *Pseudomonas aeruginosa* PA14 transcriptome, considering all genes (n = 5829), in the *ex vivo* pig lung (EVPL) model biofilm at 24 h, 48 h and 7 d post infection.** RNA was sequenced from the lung-associated biofilm at each time point from three repeats, from each of two independent pig lungs, shown by individual data points. Each time point is shown by a different shaped data point, and the individual pig lungs are shown by different colours (see key). The 95% confidence ellipses are also shown.

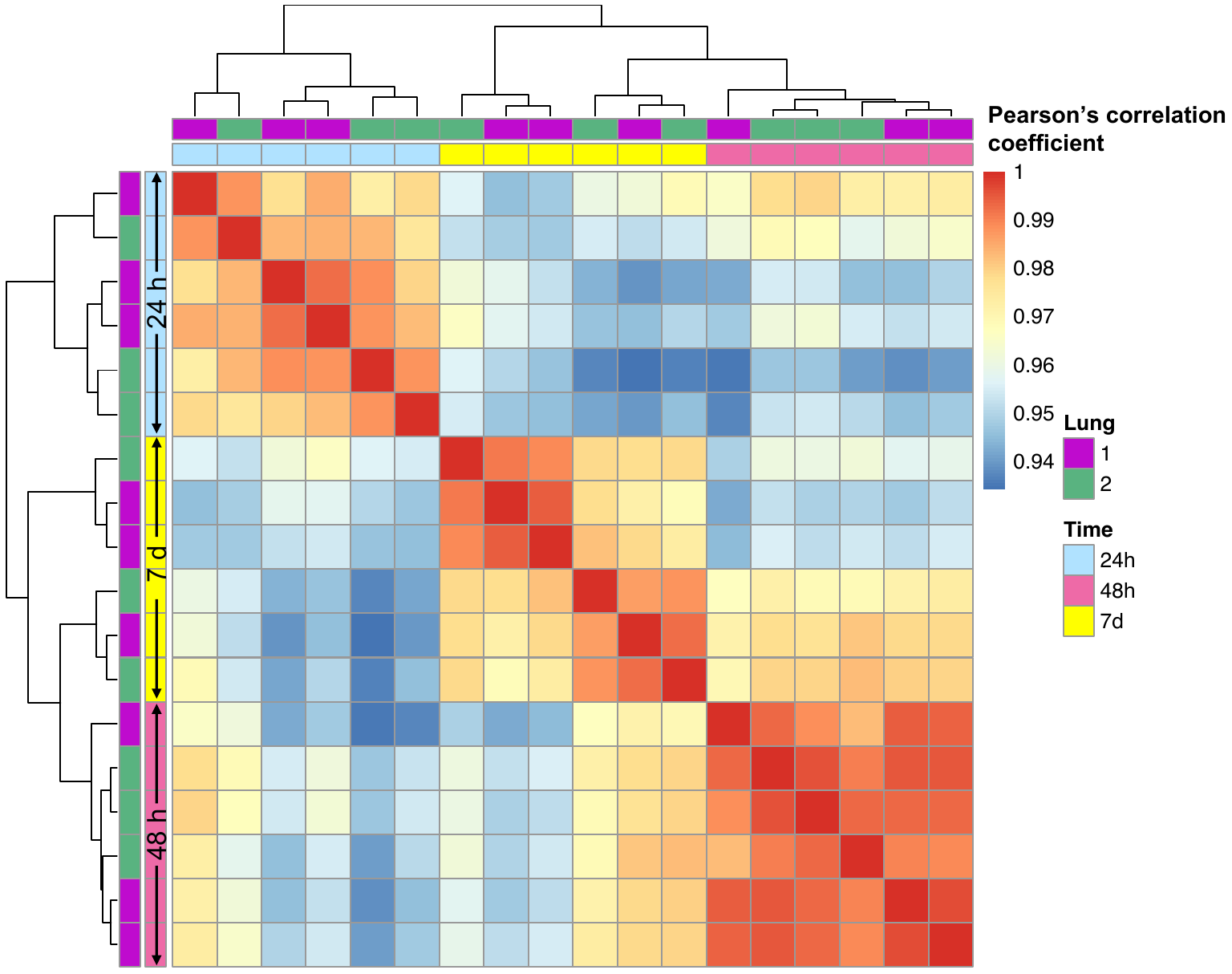
**Figure S2. Heatmap showing hierarchical clustering analysis and Pearson’s correlation coefficient values between *Pseudomonas aeruginosa* PA14 RNA sequencing data from biofilms formed on the surface of *ex vivo* pig lung (EVPL) model tissue at 24 h, 48 h and 7 d.** RNA was sequenced from the biofilm at each time point from three repeats, from each of two independent pig lungs. The independent lung each sample was grown in is shown by different colours (see key). The time point following infection RNA was extracted from is denoted by a second set of colours (see key) and labelled on the left-hand axis.

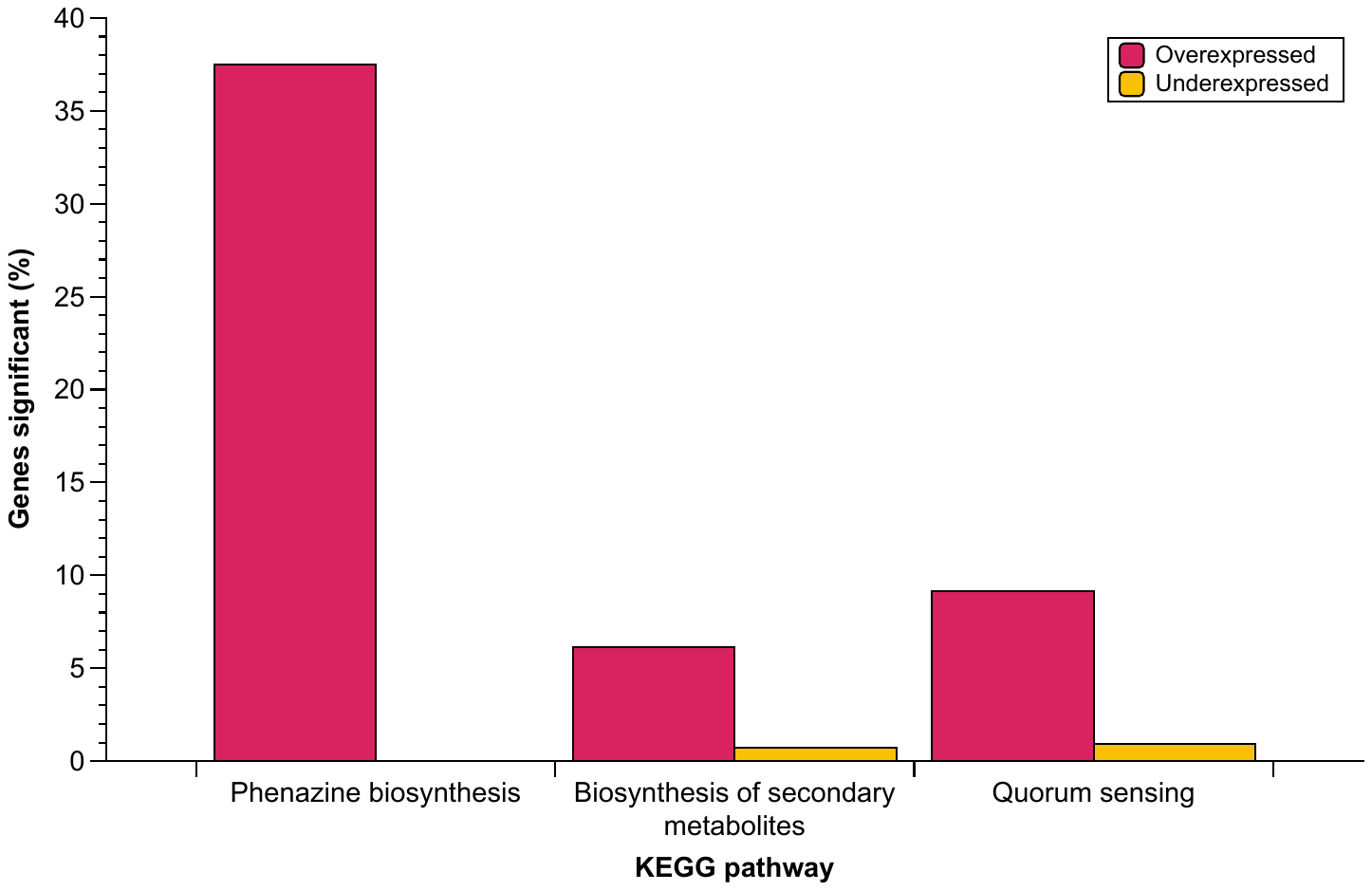
**Figure S3. Significantly enriched (*P* < 0.05) *Pseudomonas aeruginosa* PA14 Kyoto encyclopedia of genes and genomes (KEGG) pathways in the *ex vivo* pig lung-associated biofilm from 48 h to 7 d.** Each bar shows the percentage genes associated with each pathway that were significantly differentially expressed (*P* < 0.05|log_2_ fold change| ≥ 1.5). The overexpressed genes (left bar) and underexpressed genes (right bar) are show by different colour bars (see key). Three *P. aeruginosa* PA14 biofilms from each of two independent pig lungs were studied at each time point.

**Table S1. *Pseudomonas aeruginosa* PA14 expression of genes in nitrogen metabolism operons, in the *ex vivo* pig lung-associated biofilm across three time points.** Differential expression analysis was performed on three replica tissue pieces from each of two independent pig lungs at 24 h, 48 h and 7 d. Genes were considered significant if |log_2_ fold change| was ≥ 1.5, and *P* < 0.05. The log_2_ fold change values are shown in the table and were determined at the later time point compared with the earlier time point (i.e. 24 h to 48 h shows expression at 48 h compared with 24 h). The yellow fill shows genes that were significantly underexpressed at the later time point. The magenta fill represents significantly overexpressed genes, with bold and underlined font.

| **Locus tag** | **Gene name** | **Gene product** | **24 h to 48 h** | **48 h to 7 d** | **24 h to 7 d** |
| --- | --- | --- | --- | --- | --- |
| *nor* operon | | | | | |
| PA14_06810 | *norC* | Nitric-oxide reductase subunit C | -2.12 | -1.44 | -4.21 |
| PA14_06830 | *norB* | Nitric-oxide reductase subunit B | -2.38 | -1.69 | -4.40 |
| PA14_06840 |  | Denitrification protein NorD | -3.37 | -0.71 | -4.26 |
| *nir* operons | | | | | |
| PA14_06770 | *nirQ* | Regulatory protein NirQ | -1.32 | 0.13 | -1.15 |
| PA14_06790 |  | Cytochrome c oxidase subunit | -0.59 | -0.17 | -0.84 |
| PA14_06800 |  | Hypothetical protein | -0.92 | -0.30 | -1.35 |
| PA14_06650 | *nirN* | c-type cytochrome | -0.25 | **2.41** | **2.13** |
| PA14_06660 | *nirE* | Uroporphyrin-III c-methyltransferase | -0.88 | **2.51** | **1.56** |
| PA14_06670 | *nirJ* | Heme d1 biosynthesis protein NirJ | -0.38 | **2.21** | **1.78** |
| PA14_06680 | *nirH* | Hypothetical protein | -1.57 | **2.01** | 0.39 |
| PA14_06690 | *nirG* | Transcriptional regulator | -0.41 | **1.85** | 1.37 |
| PA14_06700 | *nirL* | Heme d1 biosynthesis protein NirL | -1.77 | 1.41 | -0.28 |
| PA14_06710 |  | Transcriptional regulator | -0.97 | 1.20 | 0.21 |
| PA14_06720 | *nirF* | Heme d1 biosynthesis protein NirF | -2.46 | 0.99 | -1.39 |
| PA14_06730 | *nirC* | c-type cytochrome | -1.43 | 0.59 | -0.69 |
| PA14_06740 | *nirM* | Cytochrome c-551 | -3.47 | 0.22 | -3.11 |
| PA14_06750 | *nirS* | Nitrite reductase | -2.12 | -0.43 | -2.68 |
| *nar* operons | | | | | |
| PA14_13750 | *narK1* | Nitrite extrusion protein 1 | 0.11 | -0.56 | -0.47 |
| PA14_13770 | *narK2* | Nitrite extrusion protein 2 | -0.17 | -0.07 | -0.26 |
| PA14_13780 | *narG* | Respiratory nitrate reductase alpha subunit | 0.18 | -0.69 | -0.51 |
| PA14_13800 | *narH* | Respiratory nitrate reductase beta subunit | 0.24 | -0.79 | -0.55 |
| PA14_13810 | *narJ* | Respiratory nitrate reductase delta chain | 0.24 | -0.33 | -0.13 |
| PA14_13830 | *narI* | Respiratory nitrate reductase gamma chain | 1.29 | -0.94 | 0.29 |
| PA14_13840 |  | Peptidyl-prolyl cis-trans isomerase, PpiC-type | 1.04 | -0.77 | 0.23 |
| PA14_13850 | *moaA* | Molybdenum cofactor biosynthesis protein A | 0.39 | -0.52 | -0.13 |
| PA14_13730 | *narL* | Transcriptional regulator NarL | -2.02 | 0.03 | -1.96 |
| PA14_13740 | *narX* | Two-component sensor NarX | -0.66 | -0.12 | -0.79 |
| *nos* operon | | | | | |
| PA14_20150 | *nosL* | NosL protein | -1.02 | 1.17 | 0.13 |
| PA14_20170 | *nosY* | NosY protein | 0.70 | 0.15 | 0.89 |
| PA14_20180 | *nosF* | NosF protein | 0.41 | 0.12 | 0.55 |
| PA14_20190 | *nosD* | Copper ABC transporter periplasmic substrate-binding protein | -0.71 | 0.37 | -0.29 |
| PA14_20200 | *nosZ* | Nitrous-oxide reductase | -1.18 | -0.19 | -1.48 |
| PA14_20230 | *nosR* | Regulatory protein NosR | -1.25 | -0.27 | -1.65 |
| *nap* operon | | | | | |
| PA14_30065 | *srbA* | SrbA | -0.47 | -0.09 | -0.59 |
| PA14_49210 | *napE* | Periplasmic nitrate reductase NapE | -0.47 | -0.09 | -0.59 |
| PA14_49220 | *napF* | Ferredoxin protein NapF | -1.75 | 0.32 | -1.32 |
| PA14_49230 | *napD* | NapD protein of periplasmic nitrate reductase | -00.76 | 0.23 | -0.47 |
| PA14_49250 | *napA* | Nitrate reductase catalytic subunit | -0.75 | 0.11 | -0.61 |
| PA14_49260 | *napB* | Cytochrome c-type protein NapB precursor | -0.13 | -0.14 | -0.28 |
| PA14_49270 | *napC* | Cytochrome c-type protein NapC | 0.02 | -0.10 | -0.08 |

**Table S2. *Pseudomonas aeruginosa* PA14 genes that were significantly differentially expressed in the same direction in all time contrasts in the *ex vivo* pig lung-associated biofilm.** Differential expression analysis was performed on three repeats from each of two independent pig lungs at 48 h compared with 24 h (24 h to 48 h), 7 d compared with 48 h (48 h to 7 d), and 7 d compared with 24 h (24 h to 7 d). Genes significantly differentially expressed in the same direction (i.e. overexpressed/underexpressed) in every contrast formed expression profile 1 (|log_2_ fold change| ≥ 1.5, *P* < 0.05). The log_2_ fold change value for each contrast is shown for each of these genes.

| **Locus tag** | **Gene name** | **Gene product** | **24 h to 48 h** | **48 h to 7 d** | **24 h to 7 d** |
| --- | --- | --- | --- | --- | --- |
| PA14_06830 | *norB* | Nitric-oxide reductase subunit B | -2.38 | -1.69 | -4.40 |
| PA14_22160 |  | Hypothetical protein | 1.73 | 1.61 | 3.45 |
| PA14_34740 |  | Hypothetical protein | 1.59 | 2.46 | 4.24 |
| PA14_48460 |  | Polyamine ABC transporter substrate-binding protein | 1.63 | 2.17 | 3.95 |

**Table S3. Differential expression of all 52 genes predicted to be involved in *Pseudomonas aeruginosa* PA14 antimicrobial resistance by the Comprehensive Antibiotic Resistance Database (CARD) [1].** Differential expression analysis was performed on three replica tissue pieces from each of two independent pig lungs at 24 h, 48 h and 7 d. Genes were considered significant if the |log_2_ fold change| was ≥ 1.5, and *P* < 0.05. The log_2_ fold change values are shown in the table and were determined at the later time point compared with the earlier time point (i.e. 24 h to 48 h shows expression at 48 h compared with 24 h). The model name detailed by the CARD is also shown alongside the gene locus tag and gene product.

| **Locus tag** | **Model name** | **Gene product** | **24 h to 48 h** | **48 h to 7 d** | **24 h to 7 d** |
| --- | --- | --- | --- | --- | --- |
| PA14_01940 | TriA | RND efflux membrane fusion protein | -0.30 | -0.48 | -0.80 |
| PA14_01960 | TriB | RND efflux membrane fusion protein | -1.07 | -0.20 | -1.30 |
| PA14_01970 | TriC | RND efflux transporter | -0.39 | -0.48 | -0.91 |
| PA14_05520 | MexR | Multidrug resistance operon repressor MexR | 0.31 | -0.69 | -0.37 |
| PA14_05530 | MexA | RND multidrug efflux membrane fusion protein MexA | 0.11 | -0.28 | -0.18 |
| PA14_05540 | MexB | RND multidrug efflux transporter MexB | -0.17 | -0.14 | -0.32 |
| PA14_05550 | OprM | Major intrinsic multiple antibiotic resistance efflux outer membrane protein OprM precursor | -0.97 | 0.53 | -0.38 |
| PA14_09500 | OpmD | Outer membrane protein | -1.11 | 0.74 | -0.32 |
| PA14_09520 | MexI | RND efflux transporter | -1.16 | 0.14 | -0.99 |
| PA14_09530 | MexH | RND efflux membrane fusion protein | -1.35 | 0.18 | -1.12 |
| PA14_09540 | MexG | Hypothetical protein | -1.52 | 0.32 | -1.17 |
| PA14_10470 | bcr-1 | MFS transporter | -1.04 | -1.39 | -2.49 |
| PA14_10670 | APH(3')-IIb | Aminoglycoside 3'-phosphotransferase type IIB | -0.32 | 0.55 | 0.23 |
| PA14_10790 | PDC-9 | Beta-lactamase | -0.27 | -0.09 | -0.38 |
| PA14_16280 | nalC | Transcriptional regulator | 0.17 | -0.26 | -0.09 |
| PA14_16300 | ArmR | Hypothetical protein | 0.79 | -0.10 | 0.66 |
| PA14_16790 | MexL | TetR family transcriptional regulator | -0.08 | 0.48 | -0.18 |
| PA14_16800 | MexJ | Efflux transmembrane protein | 0.47 | -0.01 | 0.46 |
| PA14_16820 | MexK | Efflux transmembrane protein | 0.05 | -0.08 | -0.03 |
| PA14_18080 | nalD | TetR family transcriptional regulator | 0.26 | -0.44 | -0.19 |
| PA14_18350 | arnA | Bifunctional UDP-glucuronic acid decarboxylase/UDP-4-amino-4-deoxy-L-arabinose formyltransferase | -1.59 | 1.22 | -0.32 |
| PA14_18760 | mexP | RND efflux membrane fusion protein | 0.36 | 0.24 | 0.62 |
| PA14_18780 | mexQ | RND efflux transporter | 0.04 | -0.18 | -0.15 |
| PA14_18790 | opmE | Outer membrane efflux protein | -0.31 | -0.03 | -0.34 |
| PA14_22760 | *Pseudomonas aeruginosa* CpxR | Two-component response regulator | -0.65 | 0.18 | -0.44 |
| PA14_31870 | MuxA | RND efflux membrane fusion protein | 0.46 | -1.41 | -0.92 |
| PA14_31890 | MuxB | RND efflux transporter | -0.58 | -0.35 | -0.95 |
| PA14_31900 | MuxC | Efflux transporter | -0.81 | -0.55 | -1.38 |
| PA14_31920 | OpmB | Outer membrane protein | -1.17 | -0.30 | -1.49 |
| PA14_32380 | OprN | Multidrug efflux outer membrane protein OprN precursor | -0.39 | 1.06 | 0.64 |
| PA14_32390 | MexF | RND multidrug efflux transporter MexF | 0.26 | 0.43 | 0.74 |
| PA14_32400 | MexE | RND multidrug efflux membrane fusion protein MexE | 0.18 | 1.05 | 1.26 |
| PA14_32410 | MexT | Transcriptional regulator MexT | -0.43 | -0.18 | -0.62 |
| PA14_32420 | MexS | Oxidoreductase | -0.36 | 0.12 | -0.23 |
| PA14_35170 | *Pseudomonas aeruginosa* soxR | Redox-sensing activator of soxS | 0.24 | 0.19 | 0.45 |
| PA14_38380 | MexZ | Transcriptional regulator | 0.51 | 0.29 | 0.83 |
| PA14_45890 | mexN | RND efflux transporter | 0.41 | -0.16 | 0.24 |
| PA14_45910 | mexM | RND efflux membrane fusion protein | 0.80 | -0.32 | 0.42 |
| PA14_46680 | PmpM | Transporter | -0.22 | -0.56 | -0.80 |
| PA14_49780 | FosA | Fosfomycin resistance protein | 0.60 | -0.11 | 0.47 |
| PA14_55170 | *Pseudomonas aeruginosa* catB7 | Chloramphenicol acetyltransferase | 0.31 | 1.04 | 1.37 |
| PA14_56880 | MexV | Membrane fusion protein | -0.42 | -0.51 | -0.93 |
| PA14_56890 | MexW | Multidrug efflux protein | 0.12 | -0.24 | -0.13 |
| PA14_59160 | CrpP | CrpP | 0.98 | -1.22 | -0.19 |
| PA14_60820 | OprJ | Outer membrane protein OprJ | -0.15 | 0.25 | 0.11 |
| PA14_60830 | MexD | Multidrug efflux RND transporter MexD | 0.32 | 0.15 | 0.50 |
| PA14_60850 | MexC | Multidrug efflux RND membrane fusion protein | 0.56 | 0.06 | 0.64 |
| PA14_60860 | Type B NfxB | Transcriptional regulator NfxB | 0.09 | 0.10 | 0.20 |
| PA14_63160 | basS | PmrB: two-component regulator system signal sensor kinase PmrB | -1.07 | 1.36 | 0.26 |
| PA14_65750 | OpmH | Outer membrane efflux protein | -1.44 | 0.29 | -1.09 |
| PA14_65990 | *Pseudomonas aeruginosa* emrE | SMR multidrug efflux transporter | 1.09 | -0.51 | 0.53 |
| PA14_72760 | OXA-50 | Beta-lactamase | 0.61 | -0.30 | 0.31 |

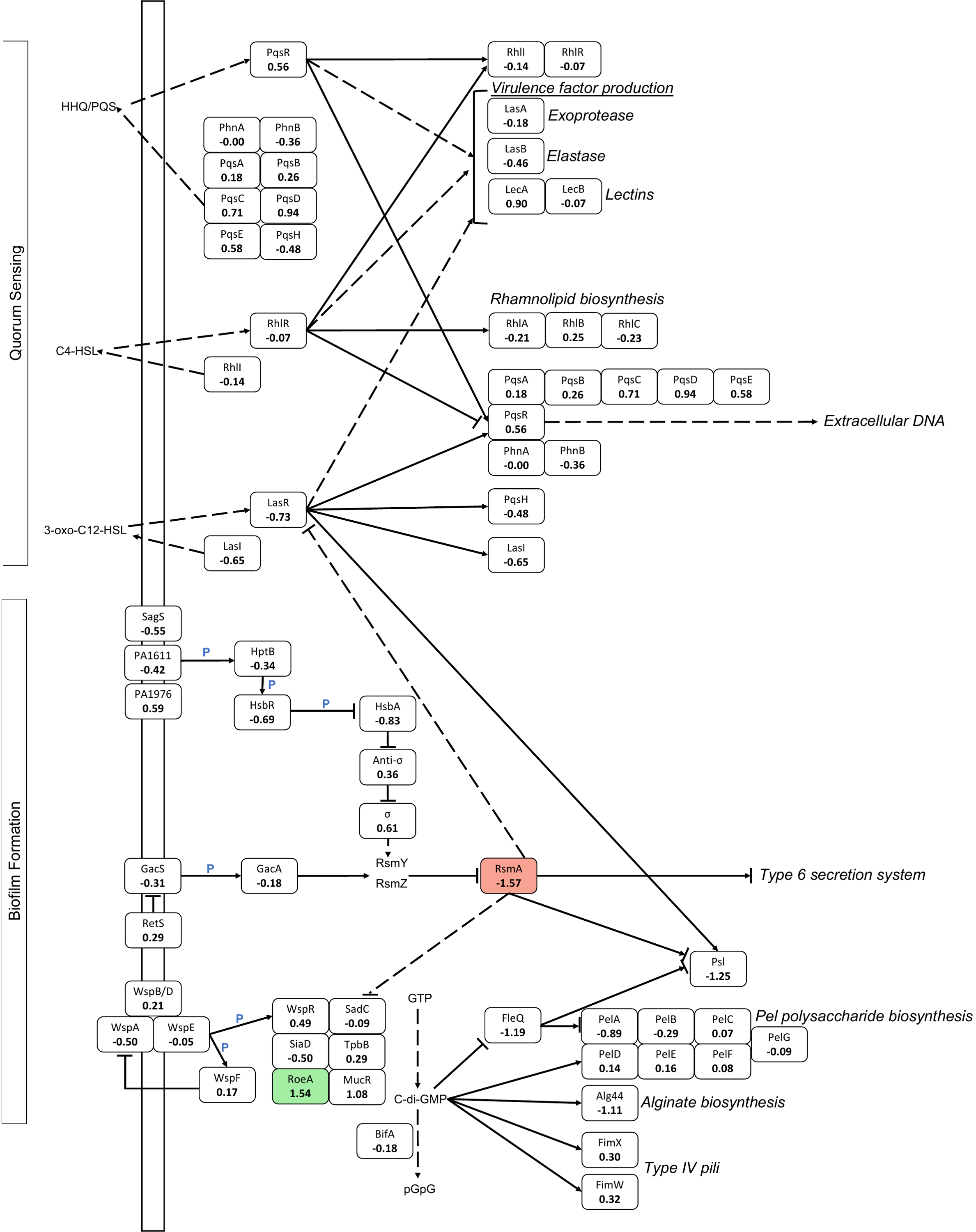
**Figure S4. Pathway map of the gene expression changes in the *Pseudomonas aeruginosa* PA14 Kyoto encyclopedia of genes and genomes (KEGG) biofilm formation and quorum sensing (QS) pathways in the *ex vivo* pig lung-associated biofilm from 48 h to 7 d.** Three repeats from each of two independent pig lungs were studied at each time point. The diagram shows all genes involved in each pathway, including associated signalling pathways and the Pel exopolysaccharide biosynthesis pathway. The log_2_ fold change value from 48 h to 7 d is shown below each gene name. The genes shown in red were significantly underexpressed, and genes shown in green were significantly overexpressed (significance: *P* < 0.05, |log_2_ fold change| ≥ 1.5). The blocked arrows represent gene activation, and the dashed arrows show indirect gene activation. The solid, blocked lines (ending with |, not an arrow) show inhibition, and the dashed, blocked lines show indirect inhibition. The blue ‘P’ represents phosphorylation.

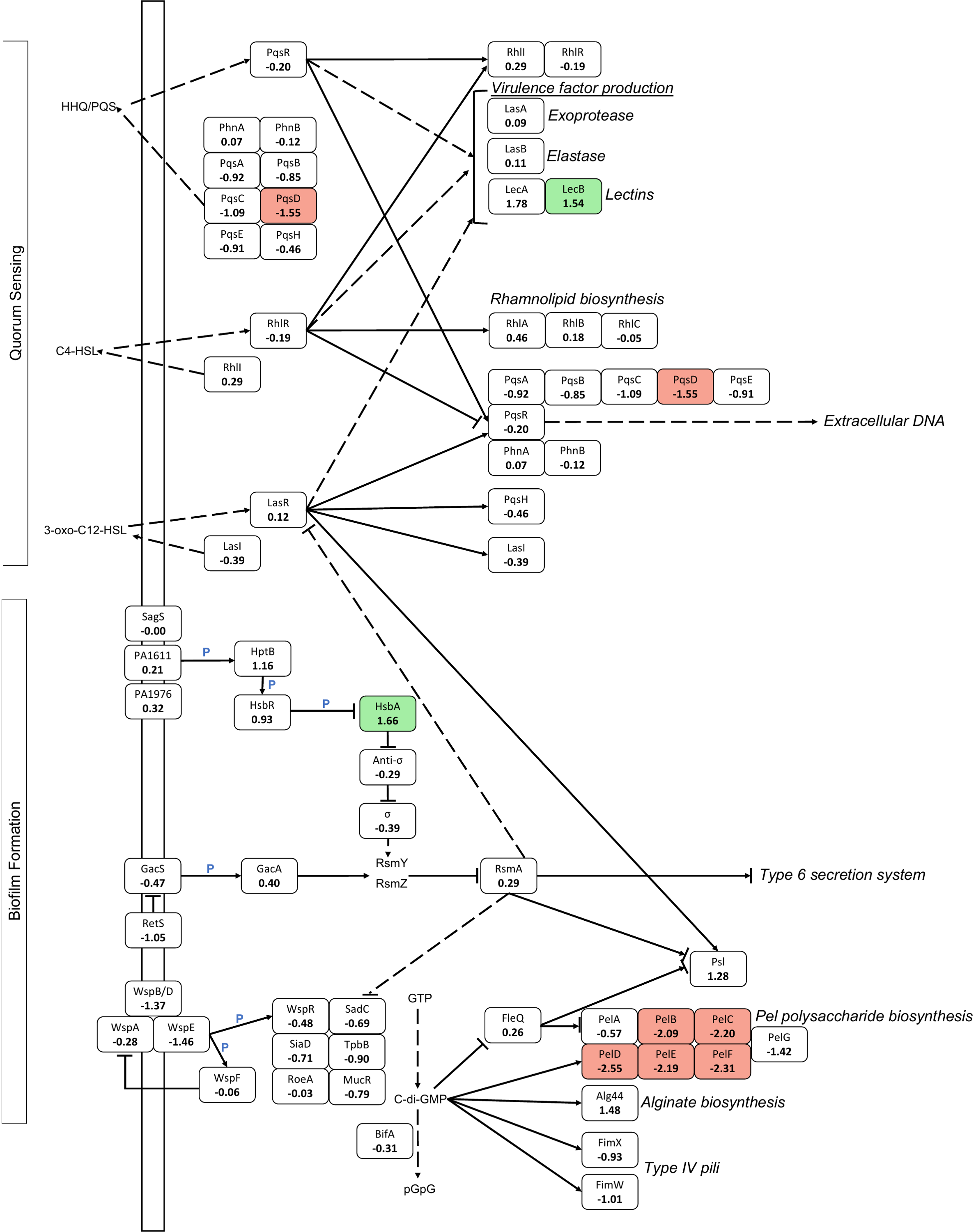
**Figure S5. Pathway map of the gene expression changes in the *Pseudomonas aeruginosa* PA14 Kyoto encyclopedia of genes and genomes (KEGG) biofilm formation and quorum sensing (QS) pathways in the *ex vivo* pig lung-associated biofilm from 24 h to 48 h.** Three repeats from each of two independent pig lungs were studied at each time point. The diagram shows all genes involved in each pathway, including associated signalling pathways and the Pel exopolysaccharide biosynthesis pathway. The log_2_ fold change value from 24 h to 48 h is shown below each gene name. The genes shown in red were significantly underexpressed, and genes shown in green were significantly overexpressed (significance: *P* < 0.05, |log_2_ fold change| ≥ 1.5). The blocked arrows represent gene activation, and the dashed arrows show indirect gene activation. The solid, blocked lines (ending with |, not an arrow) show inhibition, and the dashed, blocked lines show indirect inhibition. The blue ‘P’ represents phosphorylation.

**Figure S6. Scanning electron micrographs of uninfected *ex vivo* pig lung (EVPL) bronchiolar tissue.** The fibrous structures are consistent with the appearance of collagen fibrils and fibres under SEM. Scale bars are shown. Images were taken using a Zeiss Gemini electron microscope with an InLens detector, at 1kV, and analysed using Omero5.6 [2].
